# Evaluation of epistasis detection methods for quantitative phenotypes

**DOI:** 10.1101/2025.04.30.651312

**Authors:** Stanislav Listopad, Gauri Renjith, Qian Peng

**Affiliations:** Department of Neuroscience, The Scripps Research Institute, La Jolla, CA 92037, USA; Department of Computer Science and Engineering, University of California, San Diego, San Diego, CA 92093, USA

**Keywords:** epistasis, genetic interaction, simulation, quantitative phenotype, software benchmark

## Abstract

**Background:** Epistasis, or genetic interaction, has been increasingly recognized for its ubiquity and for its role in susceptibility to common human diseases, such as Alzheimer’s. A wide variety of epistasis detection tools are currently available with several studies comparing the performance of methods suitable for case-control data. However, there is limited understanding of how well these tools perform with quantitative phenotypes.

**Methods:** We identified six epistasis detection methods suitable for quantitative phenotype data: EpiSNP, Matrix Epistasis, MIDESP, PLINK Epistasis, QMDR, and REMMA. To evaluate these tools, we generated simulated datasets using EpiGEN. The datasets modeled various pairwise interactions between disease-associated SNPs, including dominant, multiplicative, recessive, and XOR interactions. Additionally, we assessed the BOOST and MDR algorithms on discretized (case-control) version of the datasets. These tools were then tested on the Adolescent Brain Cognitive Development (ABCD) dataset for the externalizing behavior phenotype.

**Results:** Each tool exhibited strong performance for certain interaction types, but weaker performance for others. MDR achieved the highest overall detection rate of 60%, while EpiSNP had the lowest overall detection rate of 7%. MDR and MIDESP performed best at detecting multiplicative interactions with detection rates of 54% and 41% respectively. Both MDR and MIDESP were also effective at detecting XOR interactions with detection rates of 84% and 50% respectively. PLINK Epistasis, Matrix Epistasis, and REMMA excelled at detecting dominant interactions, all achieving a 100% detection rate. On the other hand, EpiSNP was particularly effective at detecting recessive interactions with a detection rate of 66%. When analyzing the ABCD dataset, Plink Epistasis and Plink BOOST identified SNPs within the *DRD2* and *DRD4* genes, which have been previously linked to externalizing behavior.

**Conclusion:** Since no single method consistently outperforms others across all types of epistasis, and given that the specific types of epistasis present in a dataset are often unknown, it may be more effective to use multiple epistasis detection algorithms in combination to obtain comprehensive results.

## Introduction

Over a decade of genome-wide association studies (GWAS) have generated robust results for a broad range of complex traits and diseases with increasingly large sample sizes [1], [2]. While GWAS is a powerful tool for genetic mapping, most variants have shown small effects and captured a fraction of the disease heritability in general. The gap between the phenotypic variations that can be explained by genetic variants identified by GWAS and that were estimated by twin and family studies is referred to as “missing heritability” [3][4], [5]. A number of factors have been hypothesized to account for missing heritability. Genetic interaction is one such potentially critical culprit [6]. The present association studies model genetic effects of multiple variants in a linear combination (assuming additivity), while combinatorial effects beyond the linear effects are not considered. When phenotypic effects from two genetic variants deviate from the expected additive value of the individual mutations, the two variants are said to exhibit an epistatic interaction [7]. Once treated as an exception, epistasis has been increasingly recognized as ubiquitous [8], [9]. It has been shown that disease genes have a high propensity to interact with each other [10]. Additionally, the majority of genetic variants associated with complex diseases are located in non-coding regions, suggesting that they may influence traits through regulatory interactions [11].

Although epistasis has not been systematically considered in most genetic studies, evidence suggests that it contributes to susceptibility to common human diseases, such as Alzheimer’s [12], [13][14], [15]. The term epistasis may refer to biological or statistical epistasis [16], [17]. Biological epistasis involves physical interaction between two or more biological components. For instance, hair color in humans is influenced by such epistasis: the interaction between *MC1R* and *ASIP* genes is partially responsible for red hair color [18]. Meanwhile, statistical epistasis refers to the departure from additive effects of genetic variants at different loci with regard to their global contribution to the phenotype. Biological epistasis can be modeled by statistical epistasis, although statistical interaction does not always imply biological interaction. Elucidating biological interaction from statistical one is a challenging task. Improving the quality of statistical epistasis detection is nonetheless a crucial step to enable the discovery of more biological interactions. These biological interactions, may then in turn, improve our understanding of heritable traits. Given the availability of large GWAS datasets and a number of epistasis detection methods that have been developed over the years, statistical epistasis analyses may be potentially carried out at large scales. Several publications have compared the performance of epistasis detection methods primarily suitable for case-control data [19], [20], [21], [22], [23]. However, less is known about the performance of epistasis detection tools suitable for quantitative phenotypes. To this end, we conducted a survey of the commonly used epistasis detection tools, evaluating their performance on a set of simulated data, followed by testing on a real-world dataset.

For this study, we used EpiGEN to generate simulated epistasis datasets with quantitative phenotypes [24]. Given the combinatorial nature of genetic interactions, many types of epistasis theoretically exist. We modeled four major types of interactions that are most plausible: dominant, multiplicative, recessive, and exclusive-or (XOR). In the dominant model, as defined by EpiGEN, an interaction occurs if both single nucleotide polymorphisms (SNPs) each have at least one minor allele. In the multiplicative model, an interaction occurs if either SNP carries at least one minor allele, with the strength of the interaction increasing proportionally to the number of minor alleles present in the interacting SNPs. In the recessive model, an interaction is observed only if both SNPs have two minor alleles. In the XOR model, an interaction occurs only if one of the SNPs, has at least one minor allele, but not both. Note that other definitions of XOR epistasis exist, such as the one defined by Sha [25]. While the dominant, recessive, and XOR interactions are considered biologically plausible [25], [26], [27]. There is no clear evidence establishing the existence of multiplicative interactions, to the best of our knowledge. However, the multiplicative interaction is often used as an approximation for various types of interactions, especially when the exact nature of the interactions is uncertain. Throughout this publication, we use the EpiGEN-specific definitions of dominant, multiplicative, recessive, and XOR epistasis, as these terms may have alternative meanings in broader genetic contexts.

After generating the simulated data, we identified several publicly available epistasis detection tools suitable for analyzing quantitative phenotype data, including EpiSNP, Matrix Epistasis, MIDESP, PLINK Epistasis, QMDR, and REMMA [28], [29], [30], [31], [32], [33]. Our primary goal was to examine tools that could directly analyze quantitative phenotype data, which, when available, usually offers increased statistical power for detection. However, we also acknowledge that in some cases the phenotype could be discretized into a binary (case-control) value, which would allow us to utilize a wider range of analysis methods. Therefore, we also evaluated the Boolean operation-based screening and testing (BOOST) algorithm as implemented in PLINK and the multifactor dimensionality reduction (MDR) algorithm as implemented in QMDR with discretized version of our simulated phenotype [34], [35]. A complete list of tools we attempted to evaluate is provided in Table 1. While our coverage of tools is not exhaustive, we aimed to include tools that employ some of the most commonly used epistasis detection models, such as linear regression and linear mixed models, among others. We then assessed the performance of each tool in detecting dominant, recessive, multiplicative, and XOR interactions.

**Table 1:**
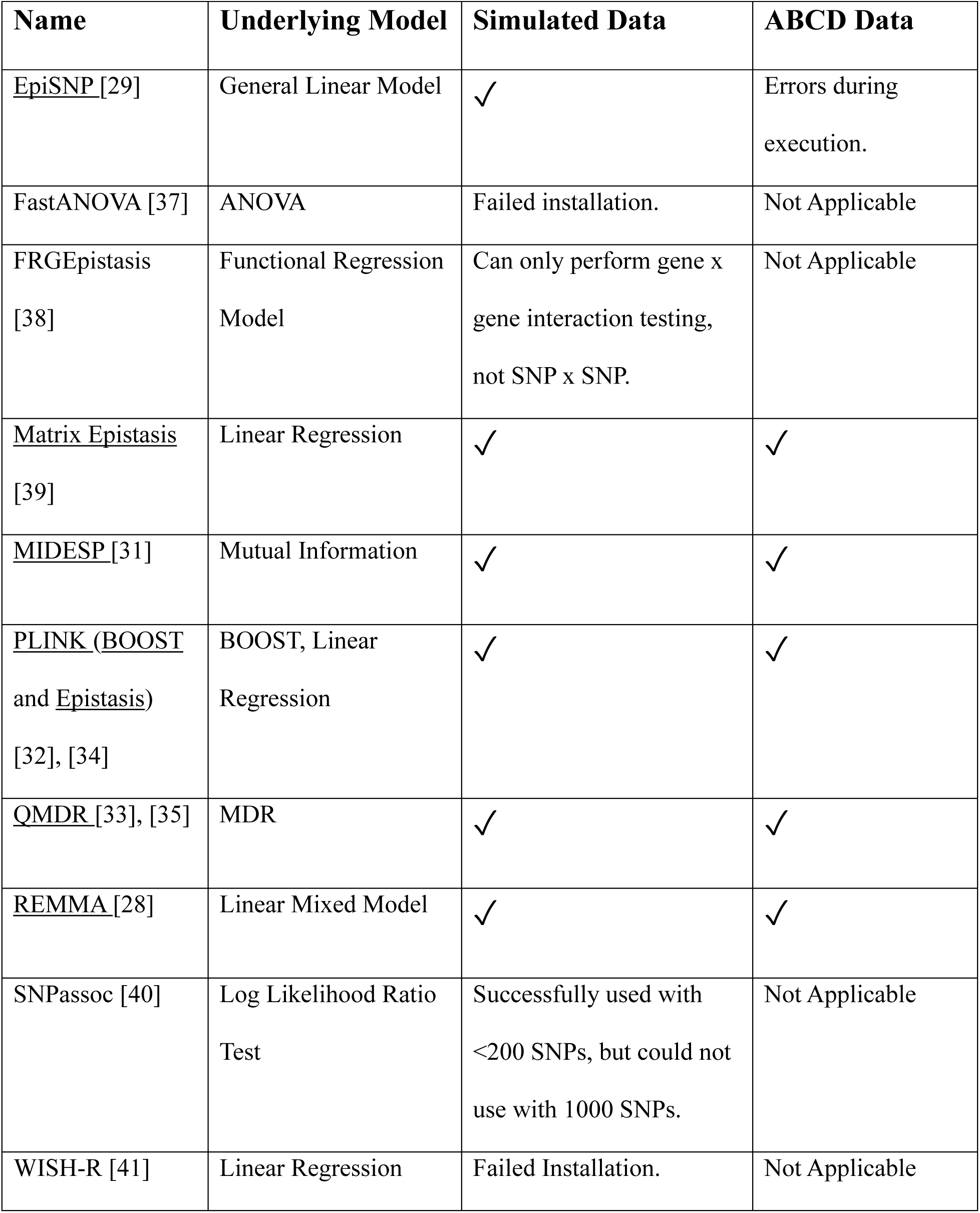
List of tools evaluated, their underlying epistasis models, and their evaluation statuses. Underlined tools were further evaluated in the study. Checkmarks (✓) indicate successful installation and usage.

| **Name** | **Underlying Model** | **Simulated Data** | **ABCD Data** |
| --- | --- | --- | --- |
| <u>EpiSNP</u> [29] | General Linear Model | ✓ | Errors during execution. |
| FastANOVA [37] | ANOVA | Failed installation. | Not Applicable |
| FRGEpistasis [38] | Functional Regression Model | Can only perform gene x gene interaction testing, not SNP x SNP. | Not Applicable |
| <u>Matrix Epistasis</u> [39] | Linear Regression | ✓ | ✓ |
| <u>MIDESP</u> [31] | Mutual Information | ✓ | ✓ |
| <u>PLINK (BOOST and Epistasis)</u> [32], [34] | BOOST, Linear Regression | ✓ | ✓ |
| <u>QMDR</u> [33], [35] | MDR | ✓ | ✓ |
| <u>REMMA</u> [28] | Linear Mixed Model | ✓ | ✓ |
| SNPassoc [40] | Log Likelihood Ratio Test | Successfully used with <200 SNPs, but could not use with 1000 SNPs. | Not Applicable |
| WISH-R [41] | Linear Regression | Failed Installation. | Not Applicable |

Epistasis can involve interactions between two or more SNPs, but this review focuses exclusively on pairwise interactions. Interaction between two SNPs is referred to as second order epistasis, while interaction between more than two SNPs is referred to as higher order epistasis. There are two categories of epistasis detection methods: exhaustive and non-exhaustive. Exhaustive algorithms test every possible combination of SNPs, which can quickly become unsustainable for higher order epistasis as the number of comparisons grows exponentially with the order of epistasis. The huge number of comparisons then results in lengthy runtime and low statistical power. Non-exhaustive epistasis detection methods mitigate these challenges by evaluating only a subset of all possible comparisons. However, there is evidence suggesting that they may perform worse than exhaustive methods [21]. For these reasons, this study focuses on 2^nd^ order epistasis detection methods.

In addition to using simulated data, we evaluated the tools in this study using the Adolescent Brain Cognitive Development (ABCD) dataset. Unlike simulated data, ABCD dataset includes population structure, individual relatedness, and multiple covariates, which add complexity to the analysis. It also contains a larger number of samples and SNPs compared to our simulated datasets. Therefore, the ABCD dataset provided a valuable opportunity to compare the performance of epistasis detection tools in a real-world context. For this study, we focused on the externalizing behavior phenotype, which is both heritable and common [36].

## Methods

### Data Generation

#### Quantitative Phenotype Datasets

A total of 40 datasets were generated using EpiGEN [24], each containing a single type of epistatic interaction: dominant, recessive, multiplicative, or XOR. Within each dataset, all interactions had the same interaction alpha [24]. Interaction alpha is a positive number that quantifies interaction strength. Definitions of dominant, recessive, and multiplicative interactions in terms of interaction alpha can be found in the EpiGEN publication. The XOR was defined as follows: if one SNP genotype is in {1, 2} (at least one minor allele) and the other SNP genotype is in {0} (no minor alleles) then there is an interaction. There are 20 multiplicative, 8 recessive, 8 dominant, and 4 XOR datasets. All interactions occurred between pairs of disease SNPs. Each dataset contained 1000 SNPs and 1000 individuals, with the data generated using chromosome 1 of the CEU HapMap3 cohort [42].

Out of the 20 multiplicative datasets, 16 contained 5 disease SNPs, and 4 contained 20 disease SNPs. The datasets with 5 disease SNPs were created by permuting the following variables: interaction alpha {1.25, 1.5, 2, 3}, number of interacting SNP pairs {1, 2}, and purity status {pure, impure}, resulting in 16 datasets. Lower interaction alphas such as 1.25 and 1.5 produce phenotype values for individuals with interacting disease SNPs that are just a few times larger than those without interacting disease SNPs, making detection challenging. Larger interaction alphas such as 2 and 3 lead to phenotype values for individuals with interacting disease SNPs to be tens of times larger than phenotype values of individuals without interacting disease SNPs, making these interactions easier to detect. Pure datasets contained no individual main effects, while impure datasets included some individual main effects [19]. The 4 datasets with 20 disease SNPs were created by permuting interaction alphas {2, 3} and purity status {pure, impure}, with each dataset containing 8 interacting disease SNP pairs. In total, the 20 multiplicative datasets contained 56 interacting SNP pairs (8 * 1 + 8 * 2 + 4 * 8 = 56).

The 8 dominant and 8 recessive datasets each contained 5 disease SNPs. These datasets were created by permuting interaction alpha values {8, 16}, the number of interacting SNP pairs {1, 2}, and purity status {pure, impure}. The 8 dominant and 8 recessive datasets contained a total of 24 interacting SNP pairs (4 * 1 + 4 * 2 + 4 * 1 + 4 * 2 = 24). The four XOR datasets each contained 20 disease SNPs with 8 interacting SNP pairs, resulting in a total of 32 interacting SNP pairs across the XOR datasets. These datasets were created by permuting interaction alphas {8, 16} and purity status {pure, impure}. Interaction alphas of 8 and 16 were chosen to ensure that the phenotypes of individuals with interacting disease SNPs were distinct from those without interacting disease SNPs. Full in-depth specifications used to generate each dataset can be found in supplemental file S1.

#### Case-Control Datasets

All of the above 40 datasets were also discretized to create an additional set of 40 case-control datasets. The discretization thresholds were selected individually for each dataset to ensure a reasonable balance of cases and controls, typically resulting in at least 1% of samples being cases. While discretizing the phenotype results in a loss of information, it enables the use of a much larger number of tools that are suitable for case-control epistasis analysis, compared to tools designed for quantitative epistasis analysis. Further details on the discretization process can be found in supplemental file S1.

#### Adolescent Brain Cognitive Development (ABCD) Dataset

The ABCD study is an ongoing project aimed at studying brain development and child health [43]. One of the more highly heritable quantitative phenotypes in the study is externalizing behavior [44], which comprises of a wide range of antisocial behaviors such as aggression, bullying, conduct problems, rule breaking, and substance use [45], [46], [47]. Due to its common occurrence among children, externalizing behavior is an ideal phenotype for study. Additionally, some genetic interactions associated with externalizing behavior, such as DRD4-DRD2 and DRD4-SLC6A4 interactions, have already been identified [48], [49], [50]. DRD4 and DRD2 are dopamine receptor genes, while SLC6A4 is a serotonin transporter gene. The ABCD study contains various externalizing scores. For this analysis, we used the child behavioral check list (CBCL) externalizing t-score, with t-score > 64 indicating clinical symptoms [51], [52].

The cohort includes 11,666 children, of which 417 had externalizing t-score > 64, after excluding 202 individuals due to missing principal components of kinship matrix. The externalizing t-scores were normalized for age and sex, and residuals were used to further adjust for population stratification (using principal components of the kinship matrix) and batch site covariates (Figure 1) [53]. For REMMA, which already accounts for population stratification, only batch site covariate was used to obtain the residuals. While some of the tools in the study, such as Matrix Epistasis, can directly account for covariates, we opted to use residuals to ensure uniformity across all tools. The discretized phenotypes were obtained by converting the residuals into case-controls with a threshold of 20.4, chosen to match the ratio of cases to controls when using the original t-score threshold of 64. Genetic data was obtained using hg19 smokescreen array as documented in the ABCD study [54], with 515,279 SNPs in the dataset. Initially, we intended to perform epistasis analysis on the entire dataset, but only PLINK was able to complete the analysis within 24 hours. Therefore, we confined the analysis to SNPs on chromosome 11, which includes 24,699 SNPs. Chromosome 11 contains the DRD4 and DRD2 genes, as well as a few other genes linked to externalizing behavior, such as NCAM1, BDNF, and TPH1 [55], [56], [57]. Limiting the analysis to a smaller number of SNPs allowed us to include all tools except EpiSNP in the comparative analysis (Table 1). EpiSNP encountered execution errors, likely due to the large dataset size. The data contained 446 SNPs with missing values. PLINK (Epistasis and BOOST), MIDESP, and REMMA were used with missing data as is. For other tools, missing SNPs were imputed with the most common non-missing values. Detailed information regarding the genetic and externalizing phenotype data can be found in the ABCD study release notes website.

**Figure 1:**
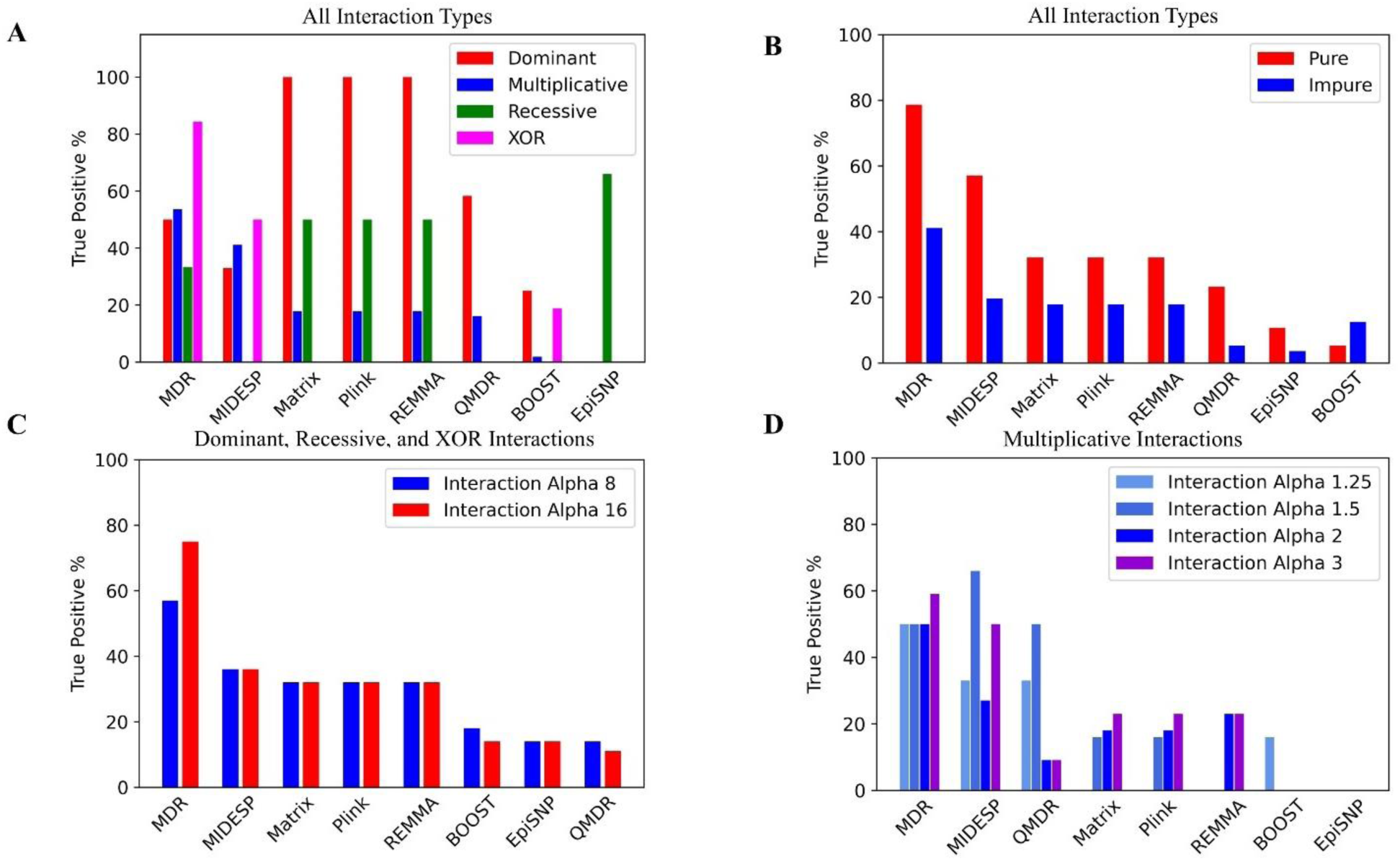
The number of successfully detected interacting SNP pairs (True Positive %) by tools and epistasis interaction types. **A. Overall performance of each** tool for each interaction type. **B**. Pure versus impure datasets. **C**. Dominant, recessive, and XOR datasets sub-grouped by interaction alpha values: 8 and 16. **D**. Multiplicative datasets sub-grouped by interaction alpha values: 1.25, 1.5, 2, and 3.

A Bonferroni correction was applied to account for multiple testing, setting the p-value significance threshold at 1.64e-10 for the analysis of 24,699 SNPs. A suggestive significance threshold of 1.64e-9 was also used. For tools that did not provide p-values, such as MIDESP, MDR, and QMDR, we retained the top 0.001% of reported results for MIDESP or the top 500 results for MDR and QMDR, based on their respective evaluation metrics (mutual information, t-statistic, and balanced accuracy). While in practice additional steps could be taken to establish the significance of the results reported by these tools, the goal of this study was to evaluate the tools based on their direct output. The interacting SNPs were then mapped onto genes using the UCSC hg19 gene annotation.

#### Epistasis Tool Selection

We began by conducting an extensive search for epistasis detection tools that could analyze quantitative phenotype data. We focused on publications offering ready-to-use software packages with available documentation. We identified ten tools that were accessible online (Table 1). Since linear regression model is widely used, we evaluated multiple tools based on this model. For other epistasis detection models, we chose the most representative tool for evaluation. After several attempts to install the tools on a personal computer and a high-performance computing (HPC) cluster, we decided to evaluate EpiSNP, Matrix Epistasis, MIDESP, PLINK Epistasis, QMDR, and REMMA. These tools were well documented, simple to install and use, and represented a diverse set of epistasis detection models. PLINK supports multiple epistasis models, and we used its linear regression model for quantitative datasets and its BOOST implementation for case-control datasets. Similarly, we applied QMDR to both quantitative and case-control datasets. We selected BOOST and QMDR for analysis of discretized data, based on previous reviews of epistasis detection algorithms [19]. For simplicity, we refer to QMDR as MDR when applied to binary phenotype data. All selected tools were well optimized and capable of analyzing > 1000 SNPs at a time.

#### Epistasis Detection Models

*Linear regression* is a linear model used to predict the value of a scalar independent variable based on dependent variable(s). It is used as the underlying model in many epistasis detection tools due to its ability to approximate various types of interactions while being easy to implement and use. The model is similar to the linear regression used in GWAS, except that it includes a term representing SNP-SNP interaction. This allows for a direct interpretation of the results [16]. Both PLINK Epistasis and Matrix Epistasis use this model to estimate interaction coefficients for each SNP pair. The key advantages of this model are its simplicity, interpretability, and fast execution. However, it has limitations, including the assumption of linear relationship between variables and its susceptibility to outliers.

*Linear mixed model* (LMM) is an extension of linear regression that allows for both fixed and random effects. LMMs offers significant advantages over traditional linear regression, particularly in genetic studies, as they can account for population stratification and individual relatedness [28]. The increased complexity of LMM, however, results in longer execution times compared to simpler models like linear regression. REMMA employs the linear mixed model in epistasis detection.

*General linear model* (GLM) is a broad class of models that includes linear regression, analysis of variance (ANOVA), and analysis of covariance (ANCOVA) [58]. Linear regression is a specific case of general linear models. EpiSNP utilizes GLMs to approximate epistasis, which is similar to, but not identical to, the approach used by PLINK and Matrix Epistasis. The advantages and disadvantages of GLMs are largely similar to those for linear regression.

*Mutual Information model* is based on information theory. It quantifies the amount of information obtained about one random variable by observing another. Mutual information (MI) has a wide variety of applications [31]. MIDESP computes the amount of mutual information shared between a subset of most promising SNP pairs and the phenotype. As a feature selection method, MI does not require any prior assumptions and can be used for both categorical and continuous data. However, its main drawback is that it can be computationally expensive to compute.

*Multifactor Dimensionality Reduction (MDR)* model is a dimensionality reduction approach for detecting interactions between independent variables influencing a dependent variable [35]. For a pair of biallelic SNPs, MDR considers nine possible genotype combinations: 0-0, 0-1, 0-2, 1-0, 1-1, 1-2, 2-0, 2-1, and 2-2 (numbers represent the number of minor alleles). For each of these combinations, a ratio of cases to controls is calculated. Genotypes where cases outnumber controls are labeled as high risk, while the opposite are labeled as low risk. Thus, a pair of SNPs, is represented as a feature with two values: low risk and high risk. A machine learning technique is then applied to identify how useful each feature is for distinguishing between cases and controls. QMDR extends this algorithm to quantitative phenotypes, by comparing mean phenotype values instead of case-control ratios. MDR transforms the data and combines it with a machine learning (ML) classifier to detect epistasis. The strength and weakness of this approach depend on the choice of ML classifier used.

#### Data Wrangling

The JSON files generated by EpiGEN had to be transformed into appropriate format for each of the selected epistasis detection tools. Several tools (MIDESP, Plink, and REMMA) use standard PLINK input files, while the rest require custom formatting.

#### Performance Evaluation

All of the selected epistasis detection tools returned lists of interacting SNP pairs. EpiSNP, Matrix Epistasis, PLINK (Epistasis and BOOST), and REMMA provided *p*-values for each SNP pair, while MIDESP provided mutual information values. QMDR provided balanced accuracy values for binary phenotype data, and t-statistic for quantitative phenotype data. We used these values to sort the SNP pairs from most to least significant. *P*-values were sorted in ascending order, while the other metrics were sorted in descending order. Each tool had a mechanism for limiting the number of output SNP pairs. For tools that used *p*-values as the sorting metric, a reporting threshold of 1e-7 was used. MIDESP reported the top 1% of hits by default, and MDR and QMDR reported up to top the 500 hits. Full details regarding the runtime configuration of each tool can be found in supplemental file S2.

The sorted lists of hits were then assessed based on three criteria: true positives, the average ranking of the true positives, and the F1 score. The average ranking of the true positives was computed using two approaches: penalized and unpenalized. In the penalized approach, for datasets with more than one interacting SNP pair, any undetected pairs were considered to be at the end of the sorted hit list. In unpenalized approach, undetected pairs were ignored. In datasets with zero true positive, the average true positive ranking was considered not applicable.

## Results

### True Positives Detected

We examined the number of true positives detected by each tool based on the type of interaction in the EpiGEN simulated data (Figure 1). The true positive percentage (%) refers to the number of successfully detected interacting SNP pairs divided by the total number of truly interacting SNP pairs present in the data. Matrix Epistasis, PLINK Epistasis, and REMMA exhibited nearly identical results, as expected, due to their similar underlying models. These tools excelled at detecting dominant epistasis interactions, identifying 100% of the true positives.

MIDESP demonstrated a drastically different performance profile compared to Matrix Epistasis, PLINK Epistasis, and REMMA. It was unable to detect recessive interactions and only identified 33% of the dominant interactions. However, MIDESP performed better at detecting multiplicative and XOR interactions, with detection rates of 41% and 50%, respectively. MDR achieved the best overall performance, with approximately 60% detection. It successfully identified 54% of all multiplicative, 50% of all dominant, 33% of all recessive, and 84% of all XOR interactions. EpiSNP performed best for recessive interactions, with a 66% detection rate, but failed to detect any other interaction types. In contrast, BOOST and QMDR exhibited generally poor performance across all interaction types, as shown in Figure 1.

### Average True Positive Ranking

While the number of successful true positive detections is important, such detections are more valuable if they are ranked highly on the sorted list of pairwise SNPs (i.e., the most significant). Otherwise, distinguishing true positives from false positives become challenging.

Therefore, for each tool, for each dataset, we calculated the unpenalized average true positive ranking (the average ranking across all interacting pairs within a single dataset) based on the interaction type (Figure 2). Datasets where no true positives were detected were excluded from this analysis. For all interaction types, except recessive, most true positives were ranked toward the top of the respective sorted lists. The mean and median unpenalized average true positive rankings for each tool and interaction type are summarized in Table 2.

**Figure 2:**
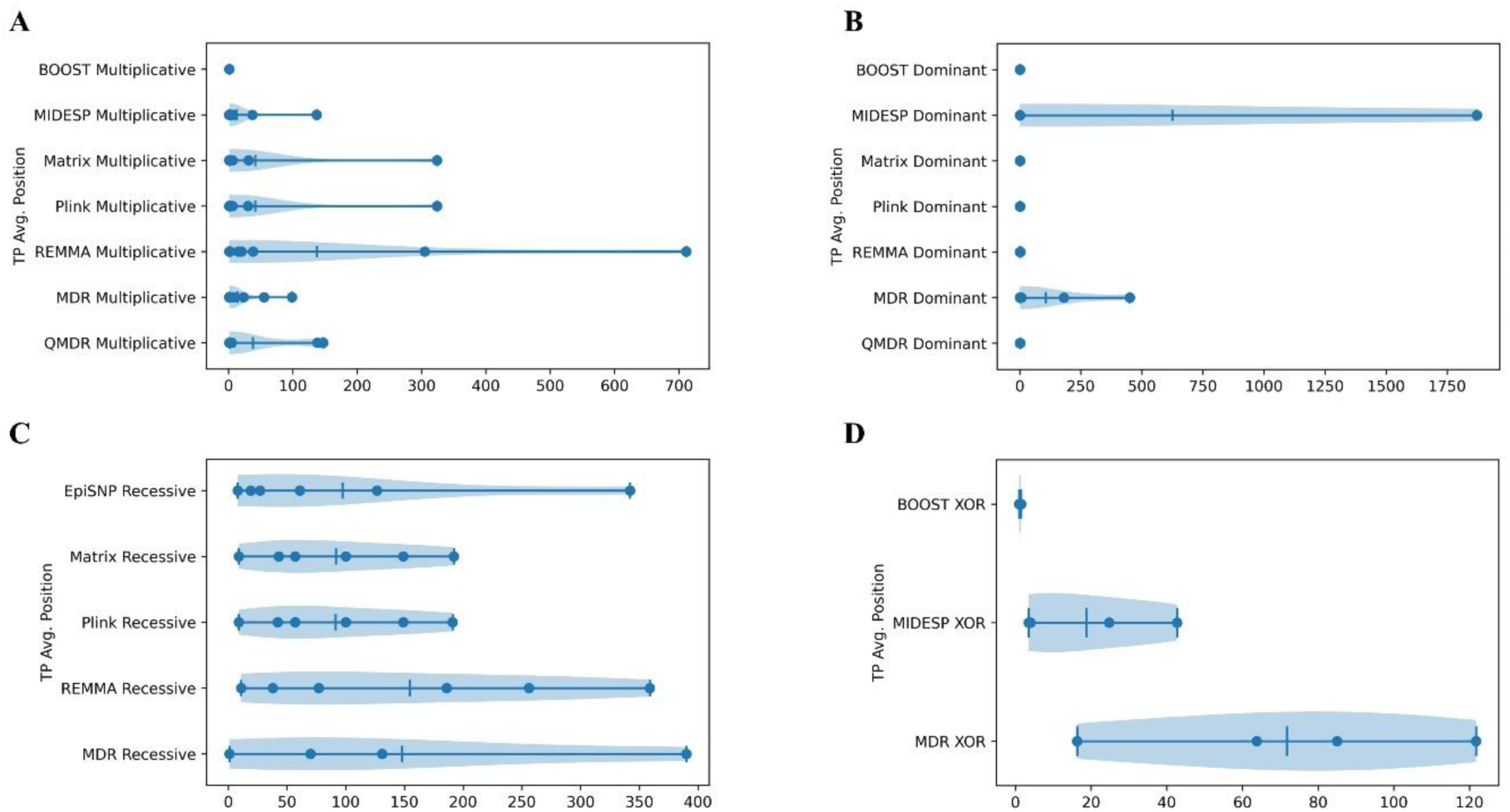
The distribution of unpenalized true positive rankings is shown as violin charts for each epistasis detection tool, categorized by interaction type: **A**. Average true positive ranking for multiplicative datasets. **B**. Average true positive ranking for dominant datasets. **C**. Average true positive ranking for recessive datasets. **D**. Average true positive ranking for XOR datasets.

**Table 2:** Summary of the mean and median unpenalized average true positive (TP) ranking for each tool and interaction type. The means and medians are listed in the following order: multiplicative (Mul.), dominant (Dom.), recessive (Rec.), and XOR. If no true positives were identified for a given interaction type, the mean/median are marked as not applicable. Values below 10 are bolded.

| Tool | Mean TP Position |  |  |  | Median TP Position |  |  |  |
| --- | --- | --- | --- | --- | --- | --- | --- | --- |
|  | Mul. | Dom. | Rec. | XOR | Mul. | Dom. | Rec. | XOR |
| MIDESP | 12.86 | 624.5 | NA | 18.75 | **2.5** | **1** | NA | 14.4 |
| Matrix Epistasis | 41.3 | **1.25** | 91.67 | NA | **3** | **1.25** | 78.5 | NA |
| PLINK Epistasis | 41.22 | **1.25** | 91.33 | NA | **3** | **1.25** | 78.5 | NA |
| PLINK BOOST | **1** | **1** | NA | 1.25 | **1** | **1** | NA | **1.25** |
| EpiSNP | NA | NA | 97.25 | NA | NA | NA | 44 | NA |
| MDR | 13.72 | 107 | 148 | 71.69 | **2.75** | **4** | 100.5 | 74.37 |
| QMDR | 37.62 | **1.25** | NA | NA | **3** | **1** | NA | NA |
| REMMA | 136.81 | **1.625** | 154.5 | NA | 17.5 | **1.5** | 131,5 | NA |

For multiplicative interactions, both MDR and MIDESP show high detection rates and high median average true positive rankings. For dominant interactions, Matrix Epistasis, PLINK Epistasis, and REMMA stand out as the top performers, exhibiting perfect detection rates and average true positive placements. For recessive interactions, EpiSNP is arguably the best tool with the highest detection rate and the best median average true positive placement. For XOR interactions, MDR demonstrates a high detection rate with poor average true positive placement. BOOST on the other hand achieves a high average true positive placement but has a poor detection rate, while MIDESP shows moderate performance in both detection rate and true positive placement.

### F1 Score

To further examine the ratio of true positives, false positives, and false negatives, we computed the F1 scores for each dataset. The mean F1 scores, averaged by interaction type for each tool, are shown in table 3. The means were computed using only datasets where at least one true positive was detected. As indicated by the low F1 values, all tools except for PLINK BOOST generated a large number of false positives (number of false negatives was negligible and can be disregarded). In this regard, PLINK BOOST is the best-performing tool.

**Table 3:** Summary of mean F1 scores for each tool, for each interaction type. Only datasets wherein at least one true positive was detected were included in the computation of the mean. Interactions for which no true positives were detected are listed as not applicable.

| Tool | Mean F1 Score |  |  |  |
| --- | --- | --- | --- | --- |
|  | Mul. | Dom. | Rec. | XOR |
| MIDESP | 0.042 | 0.0036 | NA | 0.04 |
| Matrix Epistasis | 0.038 | 0.01 | 0.002 | NA |
| PLINK Epistasis | 0.032 | 0.0099 | 0.0019 | NA |
| PLINK BOOST | 1 | 0.66 | NA | 0.3 |
| EpiSNP | NA | NA | 0.0014 | NA |
| MDR | 0.0075 | 0.004 | 0.004 | 0.026 |
| QMDR | 0.0045 | 0.0047 | NA | NA |
| REMMA | 0.0044 | 0.005 | 0.0014 | NA |

### Impact of Purity Status and Interaction Alpha

For most tools and interaction types, the detection rate was higher in pure datasets without individual main effects. The impact of interaction alpha on detection rate varied across tools. The number of interactions detected by each tool, categorized by dataset purity status and interaction alpha value, are shown in Figure 1. A detailed breakdown of each tool’s performance by interaction alpha and dataset purity status can be found in supplemental file 3.

### Runtime

We briefly evaluated the runtimes of PLINK Epistasis across a range of SNP and sample sizes. PLINK Epistasis was chosen for this evaluation because it is one of the most highly optimized tools included in the study. All analyses were executed on an HPC cluster using one node, one task, and 16 CPUs per task as assigned by the SLURM scheduler [59]. As expected, the runtimes grew approximately following an O(N^2^) order with respect to the number of SNPs (N) (Table 4). Runtimes grew linearly, O(M), with respect to the number of individuals (M). Based on these runtimes, analyzing ∼1,000,000 SNPs is feasible. In cases where analysis time becomes prohibitive, non-exhaustive algorithms might provide faster runtimes [21].

**Table 4:** PLINK Epistasis runtimes as measured on HPC (one node, one task, 16 CPUs per task) for simulated datasets with varying numbers of SNPs (1000, 10000, 100000) and individuals (1000, 5000, 10000). The estimated values are bolded and underlined.

| # of SNPs | # of Individuals | Runtime |
| --- | --- | --- |
| 1,000 | 1,000 | 0.482 seconds |
| 10,000 | 1,000 | 6.76 seconds |
| 100,000 | 1,000 | <u>7.75 minutes</u> |
| 10,000 | 5,000 | 22.2 seconds |
| 10,000 | 10,000 | 43.1 seconds |
| **<u>1,000,000</u>** | **<u>10,000</u>** | **<u>129 hours</u>** |

### ABCD Dataset

Of the six tools used to analyze ABCD data, only four provided p-values: PLINK Epistasis, PLINK Boost, Matrix Epistasis, and REMMA. The suggestive significance cutoff of 1.64e-9 was applied to the results. Using this cutoff, PLINK Epistasis reported 14 suggestively significant results, PLINK BOOST reported 178, Matrix Epistasis reported 26, and REMMA reported 2. For MIDESP, the top 0.001% of results (which contained 180 SNP pairs) was examined. For MDR and QMDR, the top 500 results were reviewed. The interacting SNPs reported by PLINK Epistasis contained rs74991942, which maps onto the DRD4 gene, while the interacting SNPs from PLINK BOOST included rs7131056, which maps onto the DRD2 gene. None of the other reported SNPs from any tool mapped to genes previously associated with externalizing behavior.

The approximate runtimes for each tool are shown in Table 5. These runtimes are provided as examples, not for direct comparison, as different hardware was used for each tool. Furthermore, REMMA can be easily parallelized to run on multiple machines using its built-in functionality, and Matrix Epistasis can also be parallelized, according to the authors [39]. MIDESP can be executed with more threads than we used in this study.

**Table 5:**
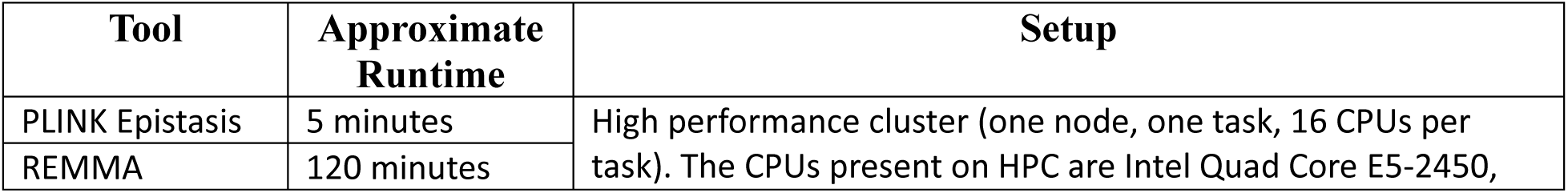
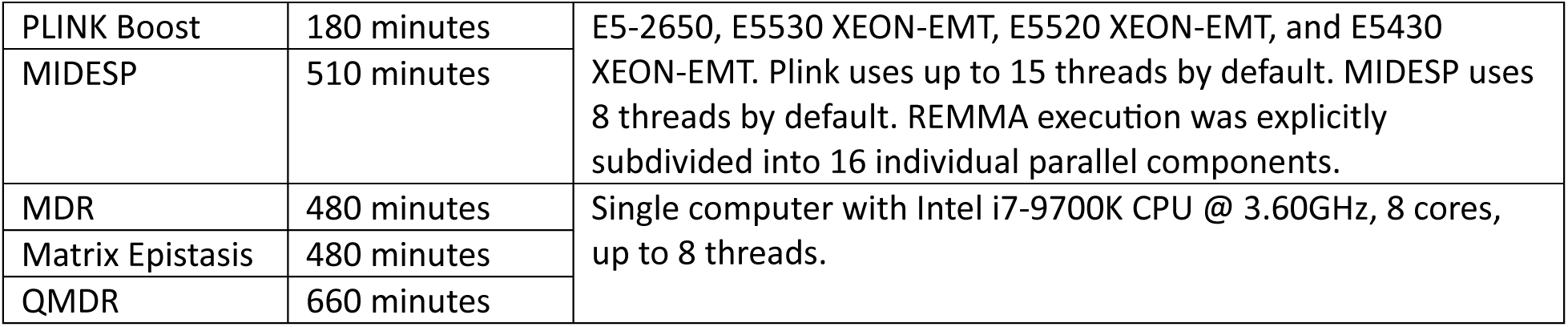
Approximate runtimes for epistasis analysis of ABCD dataset consisting of 11,666 individuals and 24,699 SNPs.

## Discussion

We conducted a survey of the epistasis detection methods for quantitative phenotypes that are suitable for large scale pairwise epistasis scans. Six tools were selected for further evaluation in this study, representing five major models for detecting epistasis: general linear models, linear mixed models, linear regression, mutual information, and multifactor dimensionality reduction. There are some recently developed tools that did not make it into the study, but are worth mentioning such as: Epi-MEIF and QuadKAST [60], [61]. Both tools are well documented and readily accessible online. While this study is not exhaustive, our focus was on providing practical recommendations to the research community, prioritizing usability and optimization over theoretical completeness.

Although the selected tools were highly optimized, processing datasets with millions of SNPs and tens of thousands of individuals remain computationally demanding. Moreover, testing trillions of pairwise interactions can quickly diminish statistical power due to the large number of multiple comparisons that must be corrected. Therefore, a filtering step may be essential to improve efficiency and maintain power. It has been shown that, despite the vast number of potential interacting gene pairs, the actual genetic interaction density is relatively low [62], and not all statistical interactions are biologically meaningful. To address this challenge, an exhaustive search method may be combined with a two-stage approach: first, a subset of genetic variants is selected in a filtering step, and then interactions are only tested among the variants within that subset. The filtering process may rely on statistical tests or biological knowledge. For instance, only genetic variants with significant main effects are subjected to interaction tests [63]. Alternatively, individual interactions could be evaluated only among variants showing significant marginal epistasis effects, which refers to the combined pairwise interaction effects between a given variant and all others [64]. The filtering stage can also incorporate biological knowledge, such as pathways linked to diseases, protein-protein interaction networks, expression quantitative trait loci (eQTL) networks, and other regulatory networks. Statistical filtering has the advantage of being discovery-driven and mostly unbiased, whereas biological filtering is hypothesis-driven, making the findings more interpretable. We have successfully used the latter strategy, coupled with a bi-clustering algorithm, to identify interacting genes associated with alcohol use disorders in an admixed population using REMMA [65].

In this study, we evaluated the performance of epistasis detection methods using simulated datasets, but we recognize that real-life datasets pose additional challenges. In real-life applications, factors such as covariates, population structure, and family structure must be accounted for [53]. Notably, of all the tools analyzed, only a few allow the inclusion of covariates directly into the epistasis analysis. Moreover, real-life datasets may contain multiple types of interactions simultaneously, whereas our simulation data focused on a specific type of interaction for each dataset.

The application of these tools to the ABCD data highlighted their advantages and limitations when handling real-world data. Tools that provide p-values for each SNP pair simplify the process of establishing significance. In contrast, other tools require additional, and sometimes time consuming steps such as permutation testing to establish significance [66]. Real datasets also tend to be much larger than our simulated ones, with many containing millions of SNPs and an increasing number of individuals. Among the tools in this study, PLINK Epistasis and REMMA are the only ones that can be easily applied to large datasets with millions of SNPs and ten thousand individuals. While REMMA is slower than PLINK, it offers the advantage of being highly parallelizable. When analyzing the ABCD data, only PLINK Epistasis and BOOST yielded the results that mapped to genes previously associated with externalizing behavior. Additionally, population stratification, individual relatedness, and other covariates such as sex and age are crucial considerations in real data analysis. REMMA stands out in this regard, as it can directly account for all of these factors. While some other tools can handle covariates, they may not fully account for complex population structures or individual relatedness directly.

Our results demonstrate that identifying interacting SNP pairs is challenging due to the large number of false positives reported by each detection method. While true positives are generally found toward the top of the sorted hit lists, this is not always the case. Consequently, further validation, including computational and biological validation, is necessary when interpreting results from epistasis detection analysis. Additionally, it is important to remember that even true statistical epistasis interactions may not always translate into meaningful biological interactions.

Our study has demonstrated that existing epistasis detection methods generally provide promising results for quantitative traits. There is however a lack of consistency in the findings across various models, which is not surprising given that the modeling frameworks differ in their respective abilities to detect specific types of epistasis. Based on our analysis, we recommend using a consensus of multiple methods to identify interacting SNP pairs in datasets with quantitative phenotypes [67]. This approach is preferable to relying on a single tool, as we cannot predict which types of epistasis may be present in real data samples *a priori*, and none of the methods evaluated in this study excelled at detecting all interaction types. However, if a single tool is preferred for convenience, PLINK Epistasis and REMMA are the most readily applicable. PLINK is best suited for datasets with low or no individual relatedness, while REMMA is uniquely suited for datasets with complex population structures and high level of individual relatedness. Notably, discretizing a quantitative phenotype (when applicable) followed by applying case-control epistasis detection methods is a viable approach.

Key Conclusions and Recommendations:

- No single tool excels at detecting all types of epistasis for quantitative phenotypes.
- Matrix Epistasis, PLINK Epistasis, and REMMA are best at detecting dominant interactions.
- MDR and MIDESP perform best at identifying multiplicative and XOR interactions.
- EpiSNP excels at detecting recessive interactions.
- PLINK Epistasis and REMMA are well suited for and readily applicable to real-world data.
- Combining tools may be the most effective approach, as the interaction types are typically unknown in real datasets.

## Supporting information

Runtime Configurations

Supplemental Results

EpiGEN Dataset Specifications

## Data Availability

All of the EpiGEN simulated data and code used to process it can be found in the following GitHub repository: https://github.com/staslist/Epistasis_Review.

## Author Biographies

**Stanislav Listopad** is a postdoc at Scripps Research under the supervision of professor Qian Peng. His research interests lie primarily in the application of software and machine learning engineering to analysis of genomic data.

**Qian Peng** is a professor of neuroscience at Scripps Research. Her research interests lie primarily in dissecting the complex genomic and epigenomic basis of addiction and related mental disorders through integrative computational approaches.

