## Supplementary material for "Evaluation of epistasis detection methods for quantitative phenotypes": Runtime Configurations

Supplementary File S2: Runtime Configurations

**EpiSNP:**

Used EPISNP1.exe with input formatted as specified in documentation.

Each dataset was represented using a ___.txt and ___chr1.dat files.

The former file contained ind_ID, father_ID, mother_ID, sex, and phenotype columns.

The latter file contained fam_ID, sex, ind_ID, and a column for every single SNP.

P-value threshold of 1e-7 was used.

**Matrix Epistasis:**

Each dataset was represented by a ____.csv and _____.pheno file.

Former contained the genotype listed for each SNP. Each column represented a single SNP. Each row represented a single individual.

Latter contained phenotypes listed for each individual in a single column.

Following R code was used to execute tool:

library("MatrixEpistasis")

compute_matrix_epistasis <- function(fname) {

fname1 = paste(fname, sep = '', '.csv')

fname2 = paste(fname, sep = '', '.pheno')

fname3 = paste(fname, sep = '', '_MatrixEpistasis.txt')

data = read.csv(fname1)

snpA = as.matrix(data)

snpB = as.matrix(data)

trait = read.csv(fname2, header = FALSE)

MatrixEpistasis_main(snpA, snpB, trait=trait, pvalThreshold=1e-7, outputFileName=fname3)

}

**MIDESP (1.2):**

Used Plink formatted file as input.

Used command as specified on MIDESP github repository to execute the tool:

java -jar MIDESP.jar -threads 70 -out Eggweight_Filtered_Pruned.epi -keep 0.25 -fdr 0.005 -apc 5000 -cont -k 30 Eggweight_Filtered_Pruned.tped Eggweight_Filtered_Pruned.tfam

java -jar MIDESP.jar -threads 70 -out Tuberculosis_Filtered_Pruned.epi -keep 0.1 -fdr 0.005 -apc 5000 Tuberculosis_Filtered_Pruned.tped Tuberculosis_Filtered_Pruned.tfam

Used all default settings (k = 30, fdr = 0.005, etc.)

-noapc flag was used, since datasets contained 1000 SNPs, and tool developers recommended not using apc with < 5000 SNPs

NOTE: it seems that ‘large’ (>5) number of significant figures in the continuous phenotype values crashes MIDESP.

**PLINK BOOST (1.90b34.2):**

Sample Command:

module load plink2/1.90b3.42

plink --file Pure_Dominant_TwoPairs_BaselineAlpha10_InteractionAlpha16_Chr1_CEU_SNP1000_IND1000_MAF005_02_converted --out Pure_Dominant_TwoPairs_BaselineAlpha10_InteractionAlpha16_Chr1_CEU_SNP1000_IND1000_MAF005_02_converted

plink --bfile Pure_Dominant_TwoPairs_BaselineAlpha10_InteractionAlpha16_Chr1_CEU_SNP1000_IND1000_MAF005_02_converted --allow-no-sex --fast-epistasis boost --epi1 1e-07 --out Pure_Dominant_TwoPairs_BaselineAlpha10_InteractionAlpha16_Chr1_CEU_SNP1000_IND1000_MAF005_02_converted_epistasis

**PLINK Epistasis (1.90b34.2):**

Sample Command:

module load plink2/1.90b3.42

plink --file Pure_Dominant_TwoPairs_BaselineAlpha10_InteractionAlpha16_Chr1_CEU_SNP1000_IND1000_MAF005_02 --out Pure_Dominant_TwoPairs_BaselineAlpha10_InteractionAlpha16_Chr1_CEU_SNP1000_IND1000_MAF005_02

plink --bfile Pure_Dominant_TwoPairs_BaselineAlpha10_InteractionAlpha16_Chr1_CEU_SNP1000_IND1000_MAF005_02 --allow-no-sex --epistasis --epi1 1e-07 --out Pure_Dominant_TwoPairs_BaselineAlpha10_InteractionAlpha16_Chr1_CEU_SNP1000_IND1000_MAF005_02_epistasis

**QMDR (3.0.2):**

Formatted data into format specified by QMDR documentation.

Used graphical interface to perform all analyses.

Loaded prepared .csv file using Load Datafile button.

Then ran analysis using Run Analysis button.

Used all default settings, except that the attribute count range was changed from 1:3 to 2:2 since we were only interested in pairwise interactions.

**MDR:**

Used QMDR tool with discretized (binary phenotype) version of our datasets.

Identical procedure to the one for QMDR.

**REMMA:**

Takes in Plink formatted files as input. REMMA was executed in five different configurations. The results from these configurations were then merged together. For any interacting SNP pair that appeared in multiple configurations the result with lowest p-value was taken. P-value threshold of 1e-7 was then used to filter the results.

REMMA can utilize five types of SNP matrices: additive (A), dominant (D), additive by additive (AxA), additive by dominant (AxD), and dominant by dominant (DxD) (Wang et al., 2020). Additionally, REMMA has three core epistasis functions: additive by additive (remma_epiAA), additive by dominant (remma_epiAD), and dominant by dominant (remma_epiDD). We combined these to create five configurations (Supplemental Table 7). In all cases we used the approximate test, instead of exact test due to runtime constraints. Code used to generate REMMA scripts can be found in our GitHub repository (<https://github.com/staslist/Epistasis_Review>) The configurations were generated based on gmat project description posted on pypi (<https://pypi.org/project/gmat/2020.4.9/>). Gmat is alternative name used by author for REMMA.

| SNP Matrices | Epistasis Test | Configuration Name |
| --- | --- | --- |
| A, AxA | remma_epiAA | aa1 |
| A, D, AxA | remma_epiAA | aa2 |
| A, D, AxA, AxD, DxD | remma_epiAA | aa3 |
| A, D, AxA, AxD, DxD | remma_epiAD | ad |
| A, D, AxA, AxD, DxD | remma_epiDD | dd |
