## Supplemental Results for "Evaluation of epistasis detection methods for quantitative phenotypes"

Supplementary File S3: Supplemental Results

Compared epistasis detection tools for quantitative phenotype.

Currently the tools are: Plink’s Epistasis, Plink’s Boost (Discretized Phenotype), Matrix Epistasis, MIDESP, MDR (Discretized Phenotype), QMDR, EpiSNP, and REMMA.

A report with results and brief discussion was generated for each tool.

The goal of this document is to compare the performance of all the included tools.

### SECTION 1 EPISTASIS TYPES, BIOLOGICAL PLAUSIBILITY

We used EpiGEN [1] to generate following interaction types:

- Multiplicative
- Dominant
- Recessive
- XOR

These epistasis interaction types are arranged in order of presumable (hypothesized) difficulty of detection.

The multiplicative, dominant, and recessive interactions are defined in EpiGEN publication.

Additionally, we implemented XOR interaction defined as follows: if one SNP genotype is in {1, 2} and the other SNP genotype is in {0} interaction alpha = alpha, otherwise interaction alpha = 1.

Total number of Epigen Datasets: 40; 20 multiplicative datasets, 8 recessive datasets, 8 dominant datasets, and 4 XOR datasets.

Total number of interacting pairs across all datasets: 112; 56 multiplicative pairs, 32 XOR pairs, 12 recessive pairs, 12 dominant pairs.

**Biological Plausibility:**

Dominant and Recessive (EpiGEN definition): several examples are provided in the Nature Education Scitable page on Epistasis [2]. Note that the definition of dominant and recessive epistasis is different within the citation from that of EPIGEN. An example of EPIGEN dominant interaction could be the one with ratio of 9:7. While an example of EPIGEN recessive interaction could be the one with ratio of 15:1.

XOR: there is limited evidence for this type of epistasis [3].

Multiplicative: unknown.

### SECTION 2 TRUE POSITIVES DETECTED

Pairs detected by tool:

- EpiSNP: 8 (7.2%)
  - Multiplicative: 0
  - Dominant: 0
  - Recessive: 8 (66%)
  - XOR: 0
- Matrix Epistasis: 28 (25%)
  - Multiplicative: 10 (17.8%)
  - Dominant: 12 (100%)
  - Recessive: 6 (50%)
  - XOR: 0 (0%)
- MDR (Discretized): 67 (59.8%)
  - Multiplicative: 30 (53.6%)
  - Dominant: 6 (50%)
  - Recessive: 4 (33.3%)
  - XOR: 27 (84.4%)
- MIDESP: 46 (41.1%)
  - Multiplicative: 23 (41.1%)
  - Dominant: 4 (33%)
  - Recessive: 0 (0%)
  - XOR: 16 (50%)
- Plink BOOST (Discretized): 10 (11.6%)
  - Multiplicative: 1 (1.78%)
  - Dominant: 3 (25%)
  - Recessive: 0 (0%)
  - XOR: 6 (18.75%)
- Plink Epistasis: 28 (25%)
  - Multiplicative: 10 (17.8%)
  - Dominant: 12 (100%)
  - Recessive: 6 (50%)
  - XOR: 0 (0%)
- QMDR: 16 (14.3%)
  - Multiplicative: 9 (16.1%)
  - Dominant: 7 (58.3%)
  - Recessive: 0
  - XOR: 0
- REMMA: 28 (25%)
  - Multiplicative: 10 (17.9%)
  - Dominant: 12 (100%)
  - Recessive: 6 (50%)
  - XOR: 0

### SECTION 3 TRUE POSITIVE AVERAGE POSITION

Now let us examine average true positive position by interaction per tool. Note that in computing the average we only account for datasets wherein at least one pair was correctly detected. Also note that these are penalized average positions, that is when a given dataset had > 1 interacting pairs and not all pairs were detected, the undetected pairs were considered to be at the “end” of the sorted list.

The detected pairs were sorted by p-value in Plink Epistasis, Matrix Epistasis, EpiSNP, and REMMA, by mutual information in MIDESP, by accuracy in MDR, and by t-statistic in QMDR.

- EpiSNP:
  - Multiplicative: 0
  - Dominant: 0
  - Recessive: 500.58
  - XOR: 0
- Matrix Epistasis:
  - Multiplicative: 247.82
  - Dominant: 1.25
  - Recessive: 414
  - XOR: NA
- MDR (Discretized):
  - Multiplicative: 110.53
  - Dominant: 179
  - Recessive: 224.37
  - XOR: 139.84
- MIDESP:
  - Multiplicative: 34.30
  - Dominant: 624.5
  - Recessive: NA
  - XOR: 72.19
- Plink BOOST (Discretized):
  - Multiplicative: 1
  - Dominant: 1.5
  - Recessive: NA
  - XOR: 2.44
- Plink Epistasis:
  - Multiplicative: 251.51
  - Dominant: 1.25
  - Recessive: 421.83
  - XOR: NA
- QMDR:
  - Multiplicative: 99.81
  - Dominant: 42.9
  - Recessive: 0
  - XOR: 0
- REMMA:
  - Multiplicative: 744.94
  - Dominant: 1.62
  - Recessive: 604.33
  - XOR: NA

Lets also examine what the average true positive positions looks like without the penalty mentioned above. Once again, we only include datasets wherein at least a single pair was detected. Let us also list the individual true positive positions (when at least one pair was detected in a dataset).

Values are listed as follows: mean; median – individual dataset average true positive positions.

- EpiSNP:
  - Multiplicative: NA
  - Dominant: NA
  - Recessive: 97.25; 44 – 8, 19, 27, 61, 126.5, 342
  - XOR: NA
- Matrix Epistasis:
  - Multiplicative: 41.3; 3 - 1, 31, 1, 1, 4, 1, 3, 324, 6
  - Dominant: 1.25; 1.25 - 1, 1, 1.5, 1.5, 1, 1, 1.5, 1.5
  - Recessive: 91.67; 78.5 - 57, 43, 192, 149, 100, 9
  - XOR: NA
- MDR (Discretized):
  - Multiplicative: 13.72; 2.75 - 2, 3, 8.5, 1, 1, 1, 1, 1, 1, 7.5, 2.5, 3.5, 55, 98.6, 10, 23
  - Dominant: 107; 4 - 1, 181, 1, 1, 7, 451
  - Recessive: 148; 100.5 - 1, 70, 131, 390
  - XOR: 71.69; 74.37 - 63.75, 121.71, 16.3, 85
- MIDESP:
  - Multiplicative: 12.86; 2.5 – 3,1,2,7,2,1,1,1,1,3,3,3,1,136.8,37,3
  - Dominant: 624.5; 1 – 1871.5, 1, 1
  - Recessive: NA
  - XOR: 18.75; 14.4 – 42.71, 24.8, 3.5, 4
- Plink BOOST (Discretized):
  - Multiplicative: 1; 1 - 1
  - Dominant: 1; 1 - 1, 1, 1
  - Recessive: NA
  - XOR: 1.25; 1.25 - 1, 1, 1.5, 1.5
- Plink Epistasis:
  - Multiplicative: 41.22; 3 – 1, 30, 1, 1, 4, 1, 3, 324, 6
  - Dominant: 1.25; 1.25 - 1, 1, 1.5, 1.5, 1, 1, 1.5, 1.5
  - Recessive: 91.33; 78.5 – 57, 42, 191, 149, 100, 9
  - XOR: NA
- QMDR:
  - Multiplicative: 37.62; 3 - 2, 1, 3, 3, 147, 5, 138, 2
  - Dominant: 1.25; 1 - 1, 2, 1, 1, 1.5, 1
  - Recessive: NA
  - XOR: NA
- REMMA:
  - Multiplicative: 136.81; 17.5 - 2, 38, 20, 711.5, 15, 1, 305, 2
  - Dominant: 1.625; 1.5 - 1, 1, 2, 3, 1, 1, 2, 2
  - Recessive: 154.5; 131.5 - 38, 77, 256, 359, 11, 186
  - XOR: NA

As we can see true positives that are detected, are often the highest ranked hit. However, many are also found outside of top 100.

Below are violin charts of unpenalized true positive positions for each tool and each epistasis interaction type.

### SECTION 4 F1 SCORE

F1 score ranges from 0 to 1 with 1 indicating perfect performance (no false positives or false negatives). F1 scores were computed for each dataset and then averaged by interaction type. When averaging the F1 scores across interaction type two computation were made (a) using all datasets (b) only using datasets where at least one true positive occurred. The (a) mean score is listed first; followed by (b) mean score, followed by – and list of all individual F1 scores per dataset.

- EpiSNP:
  - Multiplicative: 0; 0 – 0,0,0,0,0, 0,0,0,0,0, 0,0,0,0,0, 0,0,0,0,0
  - Dominant: 0; 0 – 0,0,0,0,0,0,0,0
  - Recessive: 0.0011; 0.0014 – 0, 0, 0.00067, 0.0010, 0.0010, 0.00052, 0.0034, 0.0021
  - XOR: 0; 0 – 0, 0, 0, 0
- Matrix Epistasis:
  - Multiplicative: 0.017; 0.038 – 0.0051, 0.0019, 0.023, 0.0036, 0.013, 0.1, 0.017, 0.0037, 0.18, 0,0,0,0,0,0,0,0,0,0,0
  - Dominant: 0.01; 0.01 – 0.0075, 0.0036, 0.018, 0.016, 0.0063, 0.0062, 0.013, 0.01
  - Recessive: 0.0015; 0.002 – 0,0,0.0013,0.0014, 0.0011, 0.0012, 0.0035, 0.0033
  - XOR: 0; 0 – 0,0,0,0
- MDR (Discretized):
  - Multiplicative: 0.006; 0.0075 - 0.004, 0.004, 0, 0, 0.008, 0.004, 0, 0, 0.004, 0.004, 0.004, 0.004, 0.004, 0.008, 0.008, 0.008, 0.024, 0.02, 0.0079, 0.004
  - Dominant: 0.003; 0.004 - 0, 0.004, 0, 0.004, 0.004, 0.004, 0.004, 0.004
  - Recessive: 0.002; 0.004 - 0, 0, 0, 0.004, 0.004, 0.004, 0, 0.004
  - XOR: 0.0265; 0.0265 - 0.031, 0.027, 0.024, 0.024
- MIDESP:
  - Multiplicative: 0.033; 0.042 - 0.04, 0.012, 0.067, 0.033, 0.033, 0.0047, 0.0018, 0.012, 0.1, 0.095, 0.13, 0.028, 0.014, 0.022, 0.052, 0.023, 0, 0, 0, 0
  - Dominant: 0.0013; 0.0036 - 0, 0, 0.0016, 0, 0.0031, 0.0061, 0, 0
  - Recessive: 0; 0 - 0,0,0,0, 0,0,0,0
  - XOR: 0.04; 0.04 - 0.049, 0.044, 0.046, 0.023
- Plink BOOST (Discretized):
  - Multiplicative: 0.05; 1 - 0,0,0,0,0, 0,0,0,0,0, 0,0,0,0,0, 0,0,0,0,1
  - Dominant: 0.25; 0.66 - 0,0,0.66,0.66, 0,0,0.66,0
  - Recessive: 0; 0 - 0,0,0,0, 0,0,0,0
  - XOR: 0.3; 0.3 - 0.36, 0.4, 0.22, 0.22
- Plink Epistasis:
  - Multiplicative: 0.014; 0.032 - 0.022, 0.0036, 0.013, 0.069, 0.017, 0.0036, 0.15, 0.0050, 0.0018, 0,0,0,0,0,0,0,0,0,0,0
  - Dominant: 0.0099; 0.0099 - 0.0075, 0.0035, 0.017, 0.016, 0.0061, 0.0061, 0.013, 0.0099
  - Recessive: 0.0014; 0.0019 - 0.0013, 0.0014, 0.0011, 0.0011, 0.0034, 0.0032, 0, 0
  - XOR: 0; 0 – 0,0,0,0
- QMDR:
  - Multiplicative: 0.0018; 0.0045 - 0, 0.004, 0, 0, 0, 0, 0, 0, 0.004, 0.004, 0.004, 0.004, 0.004, 0, 0.008, 0.004, 0, 0, 0, 0
  - Dominant: 0.0035; 0.0047 - 0.004, 0.004, 0, 0, 0.004, 0.004, 0.008, 0.004
  - Recessive: 0; 0 - 0, 0, 0, 0, 0, 0, 0, 0
  - XOR: 0; 0 – 0,0,0,0
- REMMA:
  - Multiplicative: 0.0018; 0.0044 - 0.0087, 0.0011, 0.0037, 0.015, 0.0028, 0.0016, 0.0018, 0.00074, 0,0,0,0, 0,0,0,0, 0,0,0,0
  - Dominant: 0.005; 0.005 - 0.0028, 0.0019, 0.0092, 0.0068, 0.0022, 0.0028, 0.0076, 0.0067
  - Recessive: 0.001; 0.0014 - 0, 0, 0.0011, 0.00095, 0.00067, 0.0011, 0.0020, 0.0024
  - XOR: 0; 0 – 0,0,0,0

### SECTION 5 MULTIPLICATIVE PURITY, INTERACTION ALPHA IMPACT

The multiplicative datasets provide a good way to examine the impact of purity and interaction alpha on rate of detection.

All average true positive positions in this section are penalized. Only multiplicative datasets are considered. Zero indicates no true positives were detected. After => is total number of true positives detected for that category divided by total number of interacting pairs in that category.

- EpiSNP:
  - Interaction Alpha 1.25: 0,0,0,0 => 0/6
  - Interaction Alpha 1.5: 0,0,0,0 => 0/6
  - Interaction Alpha 2: 0,0,0,0,0,0 => 0/22
  - Interaction Alpha 3: 0,0,0,0,0,0 => 0/22
  - Pure: 0,0,0,0,0,0,0,0,0,0 => 0/28
  - Impure: 0,0,0,0,0,0,0,0,0,0 => 0/28
- Matrix Epistasis:
  - Interaction Alpha 1.25: 0, 0, 0, 0 => 0/6
  - Interaction Alpha 1.5: 0, 0, 0, 6.5 => 1/6
  - Interaction Alpha 2: 0, 298.5, 1, 59.5, 930.87, 0 => 4/22
  - Interaction Alpha 3: 1, 0, 1, 272.5, 667.5, 0 => 5/22
  - Pure: 1, 1, 0, 0, 59.5, 272.5, 6.5, 0, 667.5, 930.87 => 8/28
  - Impure: 0, 1, 0, 0, 190.5, 0, 0, 0, 0, 0 =>2/28
- MDR (discretized):
  - Interaction Alpha 1.25: 0, 0, 1, 3.5 => 3/6
  - Interaction Alpha 1.5: 0, 0, 1, 2.5 => 3/6
  - Interaction Alpha 2: 2, 8.5, 1, 251, 249.5, 441.25 => 11/22
  - Interaction Alpha 3: 3, 251, 1, 7.5, 166.5, 378.25 => 13/22
  - Pure: 1, 1, 1, 1, 251, 7.5, 2.5, 3.5, 166.5, 249.5 => 22/28
  - Impure: 2, 3, 0, 0, 8.5, 251, 0, 0, 378.25, 441.25 => 8/28
- MIDESP:
  - Interaction Alpha 1.25: 0, 0, 1, 70.5 => 2/6
  - Interaction Alpha 1.5: 2, 0, 1, 3 => 4/6
  - Interaction Alpha 2: 3, 3, 33.5, 1, 11.5, 61.75, 0 =>6/22
  - Interaction Alpha 3: 1, 31, 1, 3, 254.25, 70.375 =>11/22
  - Pure: 1, 1, 1, 1, 11.5, 3, 3, 70.5, 254.25, 61.75 => 17/28
  - Impure: 3, 1, 2, 0, 33.5, 31, 0, 0, 70.375, 0 => 6/28
- Plink BOOST (discretized):
  - Interaction Alpha 1.25: 1, 0, 0, 0 => 1/6
  - Interaction Alpha 1.5: 0,0,0,0 =>0/6
  - Interaction Alpha 2: 0,0,0,0,0,0 =>0/22
  - Interaction Alpha 3: 0,0,0,0,0,0 =>0/22
  - Pure: 0,0,0,0,0,0,0,0,0,0 =>0/28
  - Impure: 1,0,0,0,0,0,0,0,0,0 =>1/28
- Plink Epistasis:
  - Interaction Alpha 1.25: 0, 0, 0, 0 => 0/6
  - Interaction Alpha 1.5: 0, 0, 0, 7.5 => 1/6
  - Interaction Alpha 2: 0, 293.5, 1, 61, 946.62, 0 => 4/22
  - Interaction Alpha 3: 1, 0, 1, 278.5, 673.5, 0 => 5/22
  - Pure: 1, 1, 0, 0, 61, 278.5, 7.5, 0, 673.5, 946.62 => 8/28
  - Impure: 0, 1, 0, 0, 293.5, 0, 0, 0, 0, 0, 0 => 2/28
- QMDR:
  - Interaction Alpha 1.25: 0, 0, 147, 251.5 => 2/6
  - Interaction Alpha 1.5: 0, 0, 3, 138 => 3/6
  - Interaction Alpha 2: 0, 0, 1, 253, 0, 0 => 2/22
  - Interaction Alpha 3: 2, 0, 3, 0, 0, 0 => 2/22
  - Pure: 1, 3, 3, 0, 0, 147, 253, 0, 138, 251.5 => 8/28
  - Impure: 0, 2, 0, 0, 0, 0, 0, 0, 0, 0 => 1/28
- REMMA:
  - Interaction Alpha 1.25: 0, 0, 0, 0=> 0/6
  - Interaction Alpha 1.5: 0, 0, 0, 0=> 0/6
  - Interaction Alpha 2: 0, 0, 883, 2343.12, 15, 305 => 5/22
  - Interaction Alpha 3: 0, 2, 0, 1788.875, 1, 621.5 => 5/22
  - Pure: 2343.125, 1788.875, 0, 0, 15, 1, 0, 0, 305, 621.5=> 8/28
  - Impure: 0, 0, 0, 0, 0, 2, 0, 0, 883, 0 => 2/28

### SECTION 6 DOMINANT PURITY, INTERACTION ALPHA IMPACT

All average true positive positions in this section are penalized. Only dominant datasets are considered. Zero indicates no true positives were detected.

- EpiSNP:
  - Interaction Alpha 8: 0,0,0,0 =>0/6
  - Interaction Alpha 16: 0,0,0,0 =>0/6
  - Pure: 0,0,0,0 =>0/6
  - Impure: 0,0,0,0 =>0/6
- Matrix Epistasis:
  - Interaction Alpha 8: 1, 1.5, 1, 1.5 => 6/6
  - Interaction Alpha 16: 1, 1.5, 1, 1.5 => 6/6
  - Pure: 1, 1, 1.5, 1.5 => 6/6
  - Impure: 1, 1, 1.5, 1.5 => 6/6
- MDR (discretized):
  - Interaction Alpha 8: 0, 0, 1, 254 => 2/6
  - Interaction Alpha 16: 1, 341, 1, 476 => 4/6
  - Pure: 1, 1, 254, 476 => 4/6
  - Impure: 0, 1, 0, 341 => 2/6
- MIDESP:
  - Interaction Alpha 8: 0, 1871.5, 1, 0 => 3/6
  - Interaction Alpha 16: 0, 0, 1, 0 => 1/6
  - Pure: 1, 1, 0, 0 => 2/6
  - Impure: 0, 0, 0, 1871.5 => 2/6
- Plink BOOST (discretized):
  - Interaction Alpha 8: 0, 1.5, 0, 1.5 => 2/6
  - Interaction Alpha 16: 0, 1.5, 0, 0 => 1/6
  - Pure: 0, 0, 1.5, 0 => 1/6
  - Impure: 0, 0, 1.5, 1.5 => 2/6
- Plink Epistasis:
  - Interaction Alpha 8: 1, 1.5, 1, 1.5 => 6/6
  - Interaction Alpha 16: 1, 1.5, 1, 1.5 => 6/6
  - Pure: 1, 1, 1.5, 1.5 =>6/6
  - Impure: 1, 1, 1.5, 1.5 =>6/6
- QMDR:
  - Interaction Alpha 8: 1, 0, 1, 1.5 => 4/6
  - Interaction Alpha 16: 2, 0, 1, 251 => 3/6
  - Pure: 1, 1, 1.5, 251 => 5/6
  - Impure: 1, 2, 0, 0 => 2/6
- REMMA:
  - Interaction Alpha 8: 1, 3, 1, 2 => 6/6
  - Interaction Alpha 16: 1, 2, 1, 2 => 6/6
  - Pure: 1, 1, 2, 2 => 6/6
  - Impure: 1, 2, 2, 3 => 6/6

### SECTION 7 RECESSIVE PURITY, INTERACTION ALPHA IMPACT

All average true positive positions in this section are penalized. Only recessive datasets are considered. Zero indicates no true positives were detected.

- EpiSNP:
  - Interaction Alpha 8: 0, 8, 342, 1492.5 => 4/6
  - Interaction Alpha 16: 0, 61, 126.5, 973.5 => 4/6
  - Pure: 8, 61, 126.5, 342 => 6/6
  - Impure: 0, 0, 973.5, 1492.5 => 2/6
- Matrix Epistasis:
  - Interaction Alpha 8: 0, 766, 192, 333.5 => 3/6
  - Interaction Alpha 16: 0, 734.5, 149, 309 => 3/6
  - Pure: 194, 149, 333.5, 309 => 4/6
  - Impure: 0, 0, 766, 734.5 => 2/6
- MDR (discretized):
  - Interaction Alpha 8: 0, 0, 70, 0 => 1/6
  - Interaction Alpha 16: 0, 251, 131, 445.5 => 3/6
  - Pure: 70, 131, 0, 445.5 => 3/6
  - Impure: 0, 0, 0, 251 => 1/6
- MIDESP:
  - Interaction Alpha 8: 0,0,0,0, => 0/6
  - Interaction Alpha 16: 0,0,0,0 =>0/6
  - Pure: 0,0,0, 0 =>0/6
  - Impure: 0,0,0,0 =>0/6
- Plink BOOST (discretized):
  - Interaction Alpha 8: 0,0,0,0 =>0/6
  - Interaction Alpha 16: 0,0,0,0 =>0/6
  - Pure: 0,0,0,0 =>0/6
  - Impure: 0,0,0,0 =>0/6
- Plink Epistasis:
  - Interaction Alpha 8: 0, 780.5, 191, 342 => 3/6
  - Interaction Alpha 16: 0, 753, 149, 315.5 => 3/6
  - Pure: 191, 149, 342, 315.5 => 4/6
  - Impure: 0, 0, 780.5, 753 => 2/6
- QMDR:
  - Interaction Alpha 8: 0,0,0,0 =>0/6
  - Interaction Alpha 16: 0,0,0,0 =>0/6
  - Pure: 0,0,0,0 =>0/6
  - Impure: 0,0,0,0 =>0/6
- REMMA:
  - Interaction Alpha 8: 0, 924.5, 359, 597.5=> 3/6
  - Interaction Alpha 16: 0, 256, 414.5, 1074.5 =>3/6
  - Pure: 256, 359, 414.5, 597.5 => 4/6
  - Impure: 0, 0, 1074.5, 924.5 => 2/6

### SECTION 8 XOR PURITY, INTERACTION ALPHA IMPACT

All average true positive positions in this section are penalized. Only XOR datasets are considered. Zero indicates no true positives were detected.

- EpiSNP:
  - Interaction Alpha 8: 0,0 =>0/16
  - Interaction Alpha 16: 0,0 =>0/16
  - Pure: 0,0 =>0/16
  - Impure: 0,0 =>0/16
- Matrix Epistasis:
  - Interaction Alpha 8: 0,0 =>0/16
  - Interaction Alpha 16: 0,0 =>0/16
  - Pure: 0,0 =>0/16
  - Impure: 0,0 =>0/16
- MDR (discretized):
  - Interaction Alpha 8: 169.12, 189 =>13/16
  - Interaction Alpha 16: 63.75, 137.5 =>14/16
  - Pure: 63.75, 169.12 =>15/16
  - Impure: 137.5, 189 =>12/16
- MIDESP:
  - Interaction Alpha 8: 85.37, 70.5 =>7/16
  - Interaction Alpha 16: 72, 60.875 =>9/16
  - Pure: 72, 85.37 =>13/16
  - Impure: 60.875, 70.5 =>3/16
- Plink BOOST (discretized):
  - Interaction Alpha 8: 1.87, 2.62 =>3/16
  - Interaction Alpha 16: 1.87, 3.37 =>3/16
  - Pure: 1.87, 1.87 =>2/16
  - Impure: 3.37, 2.62 =>4/16
- Plink Epistasis:
  - Interaction Alpha 8: 0,0 =>0/16
  - Interaction Alpha 16: 0,0 =>0/16
  - Pure: 0,0 =>0/16
  - Impure: 0,0 =>0/16
- QMDR:
  - Interaction Alpha 8: 0,0 =>0/16
  - Interaction Alpha 16: 0,0 =>0/16
  - Pure: 0,0 =>0/16
  - Impure: 0,0 =>0/16
- REMMA:
  - Interaction Alpha 8: 0,0 =>0/16
  - Interaction Alpha 16: 0,0 =>0/16
  - Pure: 0,0 =>0/16
  - Impure: 0,0 =>0/16

### SECTION 9 ASSEMBLY OF TOOLS

Seeing as each tool has its own strengths and weaknesses it is natural to ask whether the tools can be somehow combined in order to attain better overall performance than from any single tool.

As shown above each tool has its own strength and weaknesses.

MDR seems to be most versatile, while EpiSNP is least effective, with remaining tools performing somewhere in between.

One way to take advantage of multiple tools is by using them together. An effective scheme could be such that a detection made by at least two tools (using different underlying models) is given more weight. Especially, if this detection is near the top of the ranked list for multiple tools.

If looking for interactions of pre-determined statistical type (ex: dominant), an appropriate tool could also be selected in a targeted manner.

### SECTION 10 ADOLESCENT BRAIN COGNITIVE DEVELOPMENT DATASET

The impact of covariate adjustment on phenotype distribution is examined in supplemental figure 1.

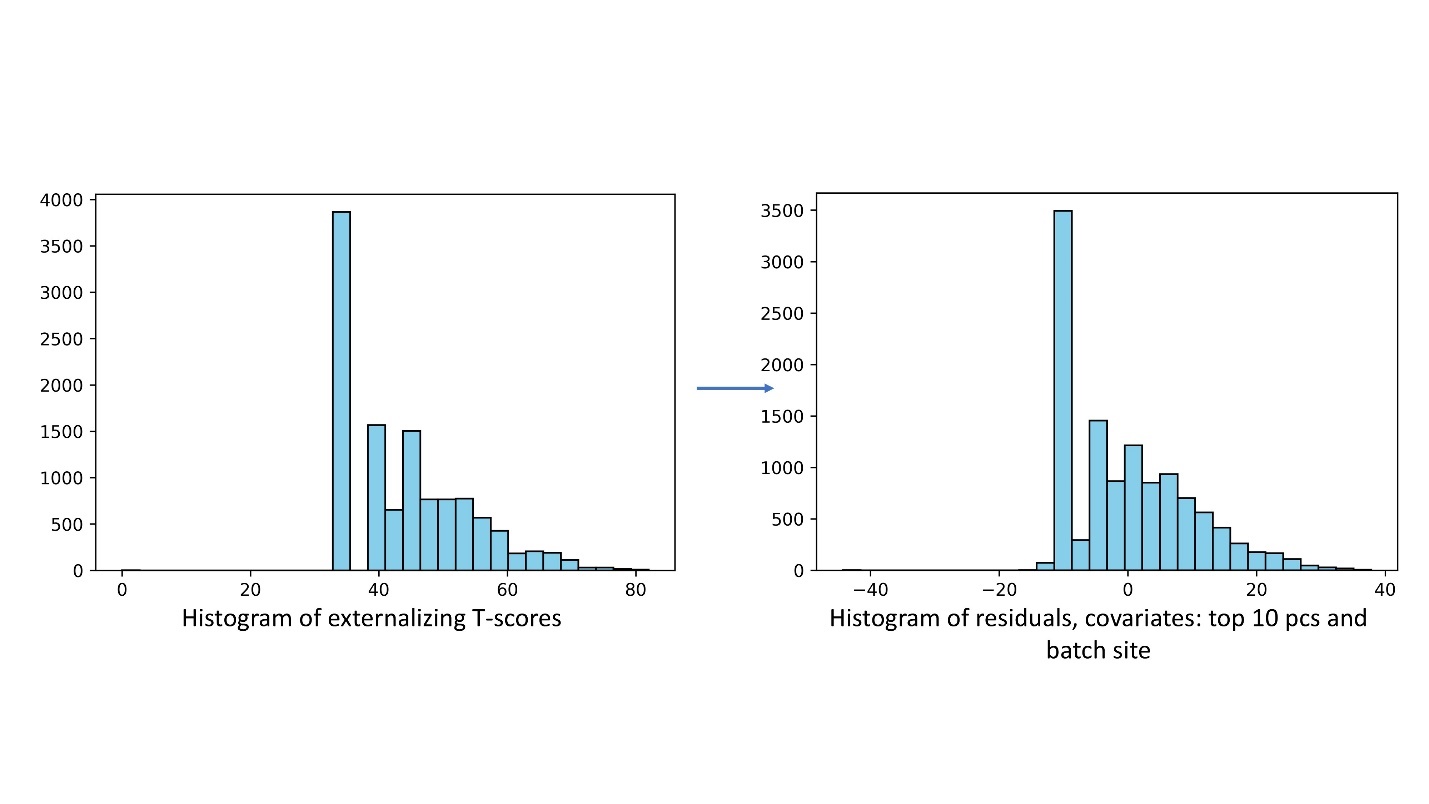

**Supplemental Figure 1: Histograms of externalizing T-scores and of residuals attained by removing the covariates**. The covariates are top ten principal components of kinship matrix and one hot encoded batch site wherein the sample (blood or saliva) was processed.
