## Supplementary material for "Evaluation of epistasis detection methods for quantitative phenotypes": EpiGEN Dataset Specifications

Supplementary File S1: EpiGEN Dataset Specifications

JSON files generated by EpiGEN for each of the datasets specified below can be found in the following GitHub repository: <https://github.com/staslist/Epistasis_Review>

The XML files specifying the disease SNPs and their interactions can be also be found in the same repository.

Multiplicative dataset specifications:

- Epistasis Type: Pure, Impure
- Interaction Order: 2^nd^
  - Number of Interacting (non-overlapping) Pairs: 1, 2, and 8
  - There are 8 datasets with 1 interacting pair, 8 datasets with 2 interacting pairs, and 4 datasets with 8 interacting pairs
- Interaction Models: multiplicative (all pairs are same interaction model with same interaction alpha)
- Number of SNPs: 1000
- Number of Individuals: 1000
- Minor Allele Frequency Range: 0.05 - 0.2
- Chromosome: 1
- Population: CEU
- Phenotype: quantitative
- Baseline Alpha: 10
- Marginal Model: additive + recessive + dominant (mixed for impure, all recessive for pure)
- Number of Disease SNPs:
  - 5 such that there are 1 or 2 interacting pairs
  - 20 such that there are 8 interacting pairs
- For datasets with 5 disease SNPs (16 datasets):
  - Interaction Alpha: 1.25, 1.5, 2, 3
    - Four datasets with interaction alpha 1.25
    - Four datasets with interaction alpha 1.5
    - Four datasets with interaction alpha 2
    - Four datasets with interaction alpha 3
  - Marginal Alpha:
    - Pure: recessive model with alpha 1.0 (no impact)
      - Ten datasets are pure with no individual main effects
    - Impure:
      - Two SNPs are recessive marginal model with alpha 0.5
      - Two SNPs are additive marginal model with alpha 2
      - One SNPs is dominant marginal model with alpha 3
      - Ten datasets are impure with above-stated individual main effects
- For datasets with 20 disease SNPs (4 datasets):
  - Interaction Alpha:
    - Two datasets with interaction alpha 2
    - Two datasets with interaction alpha 3
  - Marginal Alpha:
    - Pure: recessive model with alpha 1.0 (no impact)
      - Two datasets are pure with no individual main effects
    - Impure:
      - Ten SNPs are recessive marginal model with alphas: 1.0, 1.1, 1.2, 1.3, 1.4, 1.5, 1.6, 1.7, 1.8, 1.9
      - Five SNPs are additive marginal model with alphas: 0.8, 0.85, 0.9, 0.95, 1.0
      - Five SNPs are dominant marginal model with alpha: 1.25
      - Two datasets are impure with above-stated individual main effects

Resulting multiplicative datasets:

Pure_Multiplicative_TwoPairs_BaselineAlpha10_InteractionAlpha125_Chr1_CEU_SNP1000_IND1000_MAF005_02

Pure_Multiplicative_TwoPairs_BaselineAlpha10_InteractionAlpha15_Chr1_CEU_SNP1000_IND1000_MAF005_02

Pure_Multiplicative_TwoPairs_BaselineAlpha10_InteractionAlpha2_Chr1_CEU_SNP1000_IND1000_MAF005_02

Pure_Multiplicative_TwoPairs_BaselineAlpha10_InteractionAlpha3_Chr1_CEU_SNP1000_IND1000_MAF005_02

Pure_Multiplicative_OnePair_BaselineAlpha10_InteractionAlpha125_Chr1_CEU_SNP1000_IND1000_MAF005_02

Pure_Multiplicative_OnePair_BaselineAlpha10_InteractionAlpha15_Chr1_CEU_SNP1000_IND1000_MAF005_02

Pure_Multiplicative_OnePair_BaselineAlpha10_InteractionAlpha2_Chr1_CEU_SNP1000_IND1000_MAF005_02

Pure_Multiplicative_OnePair_BaselineAlpha10_InteractionAlpha3_Chr1_CEU_SNP1000_IND1000_MAF005_02

Pure_Multiplicative_EightPairs_BaselineAlpha10_InteractionAlpha2_Chr1_CEU_SNP1000_IND1000_MAF005_02

Pure_Multiplicative_EightPairs_BaselineAlpha10_InteractionAlpha3_Chr1_CEU_SNP1000_IND1000_MAF005_02

Impure_Multiplicative_TwoPairs_BaselineAlpha10_InteractionAlpha125_Chr1_CEU_SNP1000_IND1000_MAF005_02

Impure_Multiplicative_TwoPairs_BaselineAlpha10_InteractionAlpha15_Chr1_CEU_SNP1000_IND1000_MAF005_02

Impure_Multiplicative_TwoPairs_BaselineAlpha10_InteractionAlpha2_Chr1_CEU_SNP1000_IND1000_MAF005_02

Impure_Multiplicative_TwoPairs_BaselineAlpha10_InteractionAlpha3_Chr1_CEU_SNP1000_IND1000_MAF005_02

Impure_Multiplicative_OnePair_BaselineAlpha10_InteractionAlpha125_Chr1_CEU_SNP1000_IND1000_MAF005_02

Impure_Multiplicative_OnePair_BaselineAlpha10_InteractionAlpha15_Chr1_CEU_SNP1000_IND1000_MAF005_02

Impure_Multiplicative_OnePair_BaselineAlpha10_InteractionAlpha2_Chr1_CEU_SNP1000_IND1000_MAF005_02

Impure_Multiplicative_OnePair_BaselineAlpha10_InteractionAlpha3_Chr1_CEU_SNP1000_IND1000_MAF005_02

Impure_Multiplicative_EightPairs_BaselineAlpha10_InteractionAlpha2_Chr1_CEU_SNP1000_IND1000_MAF005_02

Impure_Multiplicative_EightPairs_BaselineAlpha10_InteractionAlpha3_Chr1_CEU_SNP1000_IND1000_MAF005_02

Recessive and dominant datasets:

- Epistasis Type: Pure, Impure
- Interaction Order: 2^nd^
  - Number of Interacting (non-overlapping) Pairs: 1, 2 pairs
  - There are 8 datasets with 1 interacting pair and 8 datasets with 2 interacting pairs
    - Four 1 pair datasets and four 2 pair datasets are dominant interaction
    - Four 1 pair datasets and four 2 pair datasets are recessive interaction
- Interaction Models: recessive, dominant (all pairs are same interaction model with same interaction alpha)
- Number of SNPs: 1000
- Number of Individuals: 1000
- Minor Allele Frequency Range: 0.25-0.4 for recessive datasets, 0.05-0.2 for dominant datasets
- Chromosome: 1
- Population: CEU
- Phenotype: quantitative
- Baseline Alpha: 10
- Marginal Model: additive + recessive + dominant (mixed for impure, all recessive for pure)
- Number of Disease SNPs: 5
- Interaction Alpha 2, 3
  - Eight datasets with interaction alpha 2
  - Eight datasets with interaction alpha 3
- Marginal Alpha:
  - Pure: recessive model with alpha 1.0 (no impact)
    - Eight datasets are pure with no individual main effects
  - Impure:
    - Two SNPs are recessive marginal model with alpha 0.5
    - Two SNPs are additive marginal model with alpha 2
    - One SNPs is dominant marginal model with alpha 3
    - Ten datasets are impure with above-stated individual main effects

Resulting recessive and dominant datasets:

Pure_Recessive_OnePair_BaselineAlpha10_InteractionAlpha8_Chr1_CEU_SNP1000_IND1000_MAF025_04

Pure_Recessive_OnePair_BaselineAlpha10_InteractionAlpha16_Chr1_CEU_SNP1000_IND1000_MAF025_04

Pure_Recessive_TwoPairs_BaselineAlpha10_InteractionAlpha16_Chr1_CEU_SNP1000_IND1000_MAF025_04

Pure_Recessive_TwoPairs_BaselineAlpha10_InteractionAlpha8_Chr1_CEU_SNP1000_IND1000_MAF025_04

Impure_Recessive_TwoPairs_BaselineAlpha10_InteractionAlpha8_Chr1_CEU_SNP1000_IND1000_MAF025_04

Impure_Recessive_TwoPairs_BaselineAlpha10_InteractionAlpha16_Chr1_CEU_SNP1000_IND1000_MAF025_04

Impure_Recessive_OnePair_BaselineAlpha10_InteractionAlpha16_Chr1_CEU_SNP1000_IND1000_MAF025_04

Impure_Recessive_OnePair_BaselineAlpha10_InteractionAlpha8_Chr1_CEU_SNP1000_IND1000_MAF025_04

Pure_Dominant_OnePair_BaselineAlpha10_InteractionAlpha8_Chr1_CEU_SNP1000_IND1000_MAF005_02

Pure_Dominant_OnePair_BaselineAlpha10_InteractionAlpha16_Chr1_CEU_SNP1000_IND1000_MAF005_02

Pure_Dominant_TwoPairs_BaselineAlpha10_InteractionAlpha8_Chr1_CEU_SNP1000_IND1000_MAF005_02

Pure_Dominant_TwoPairs_BaselineAlpha10_InteractionAlpha16_Chr1_CEU_SNP1000_IND1000_MAF005_02

Impure_Dominant_OnePair_BaselineAlpha10_InteractionAlpha8_Chr1_CEU_SNP1000_IND1000_MAF005_02

Impure_Dominant_OnePair_BaselineAlpha10_InteractionAlpha16_Chr1_CEU_SNP1000_IND1000_MAF005_02

Impure_Dominant_TwoPairs_BaselineAlpha10_InteractionAlpha8_Chr1_CEU_SNP1000_IND1000_MAF005_02

Impure_Dominant_TwoPairs_BaselineAlpha10_InteractionAlpha16_Chr1_CEU_SNP1000_IND1000_MAF005_02

XOR Datasets:

- Epistasis Type: Pure, Impure
- Interaction Order: 2^nd^
  - Number of Interacting (non-overlapping) Pairs: 8 pairs
  - There are 4 datasets with 8 interacting pairs
- Interaction Models: XOR (all pairs are same interaction model with same interaction alpha)
- Number of SNPs: 1000
- Number of Individuals: 1000
- Minor Allele Frequency Range: 0.05-0.2 for dominant datasets
- Chromosome: 1
- Population: CEU
- Phenotype: quantitative
- Baseline Alpha: 10
- Marginal Model: additive + recessive + dominant (mixed for impure, all recessive for pure)
- Number of Disease SNPs: 20
- Interaction Alpha: 8, 16
  - Two datasets with interaction alpha 8
  - Two datasets with interaction alpha 16
- Marginal Alpha:
  - Marginal Alpha:
    - Pure: recessive model with alpha 1.0 (no impact)
      - Two datasets are pure with no individual main effects
    - Impure:
      - Ten SNPs are recessive marginal model with alphas: 1.0, 1.1, 1.2, 1.3, 1.4, 1.5, 1.6, 1.7, 1.8, 1.9
      - Five SNPs are additive marginal model with alphas: 0.8, 0.85, 0.9, 0.95, 1.0
      - Five SNPs are dominant marginal model with alpha: 1.25
      - Two datasets are impure with above-stated individual main effects

Resulting XOR datasets:

Impure_XOR_EightPairs_BaselineAlpha10_InteractionAlpha8_Chr1_CEU_SNP1000_IND1000_MAF005_02

Impure_XOR_EightPairs_BaselineAlpha10_InteractionAlpha16_Chr1_CEU_SNP1000_IND1000_MAF005_02

Pure_XOR_EightPairs_BaselineAlpha10_InteractionAlpha8_Chr1_CEU_SNP1000_IND1000_MAF005_02

Pure_XOR_EightPairs_BaselineAlpha10_InteractionAlpha16_Chr1_CEU_SNP1000_IND1000_MAF005_02

Phenotype Binarization Thresholds:

The threshold is selected such that if there is an effective interaction between at least one disease SNP pair the phenotype is case.

Multiplicative Interaction Alpha 1.25: 15

Multiplicative Interaction Alpha 1.5: 22

Multiplicative Interaction Alpha 2: 40

Multiplicative Interaction Alpha 3: 90

Recessive Interaction Alpha 8: 30

Recessive Interaction Alpha 16: 60

Dominant Interaction Alpha 8: 30

Dominant Interaction Alpha 16: 60

XOR Interaction Alpha 8: 30

XOR Interaction Alpha 16: 60
